# miR-486 is an epigenetic modulator of Duchenne muscular dystrophy pathologies

**DOI:** 10.1101/2021.06.14.448387

**Authors:** Rylie M. Hightower, Adrienne Samani, Andrea L. Reid, Katherine G. English, Michael A. Lopez, J. Scott Doyle, Michael J. Conklin, David A. Schneider, Marcas M. Bamman, Jeffrey J. Widrick, David K. Crossman, Min Xie, David Jee, Eric C. Lai, Matthew S. Alexander

## Abstract

Duchenne muscular dystrophy (DMD) is an X-linked progressive muscle disorder resulting in muscle weakness and cardiomyopathy. MicroRNAs have been shown to play essential roles in muscle development, metabolism, and disease pathologies. We demonstrated that miR-486 expression is reduced in DMD muscles and its expression levels correlate with dystrophic disease severity. *MicroRNA-486* knockout mice developed disrupted myofiber architecture, decreased myofiber size, decreased locomotor activity, increased cardiac fibrosis, and metabolic defects that were exacerbated on the dystrophic *mdx^5cv^* background. We integrated RNA-sequencing and chimeric eCLIP-sequencing data to identify direct *in vivo* targets of miR-486 and associated dysregulated gene signatures in skeletal muscle. In comparison to our DMD mouse muscle transcriptomes, we identified several of these miR-486 muscle targets including known modulators of dystrophinopathy disease symptoms. Together, our studies identify miR-486 as a driver of muscle remodeling in DMD, a useful biomarker for dystrophic disease progression, and highlight chimeric eCLIP-sequencing as a useful tool to identify direct *in vivo* microRNA target transcripts.

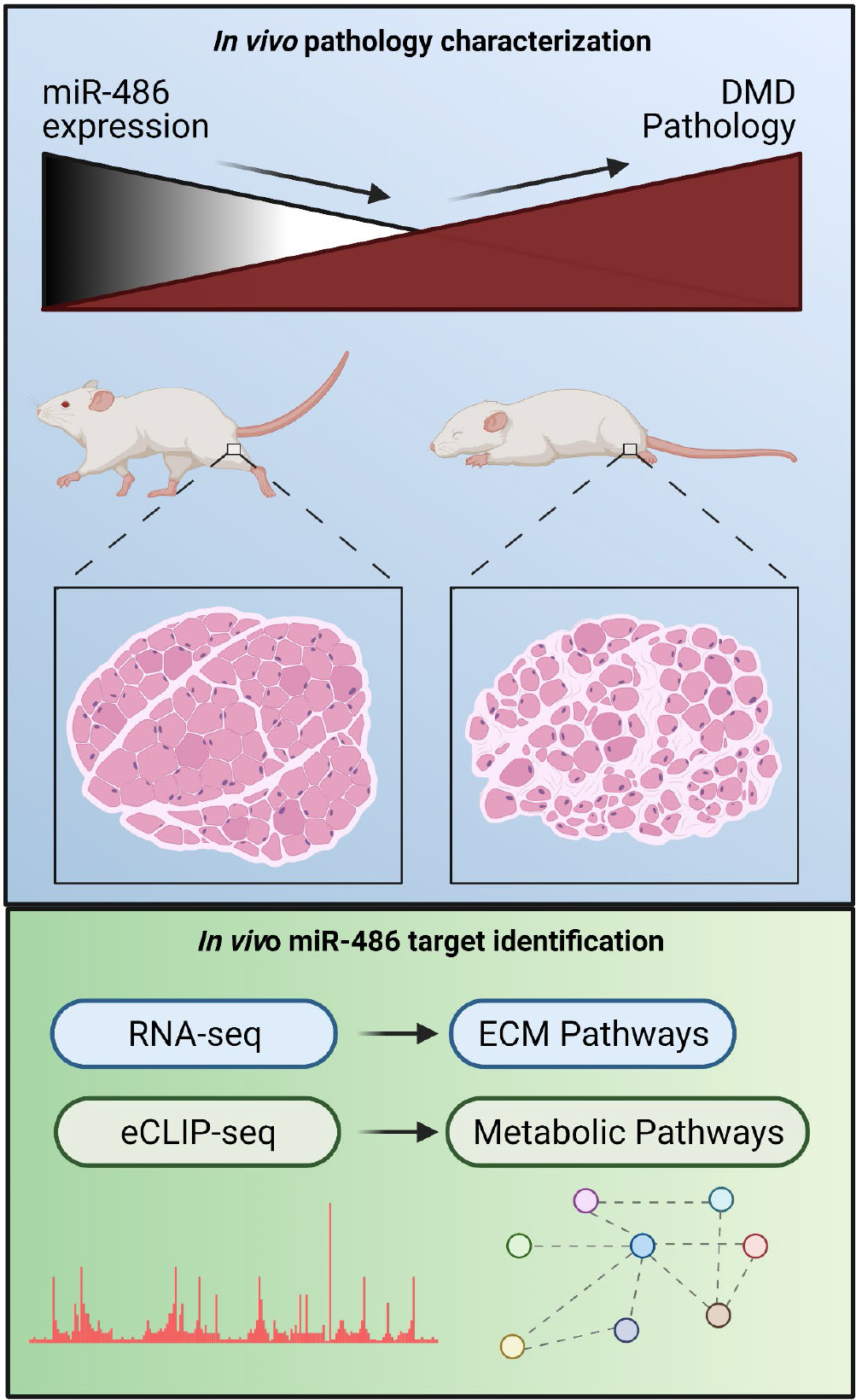

## Main

Skeletal muscle is a remarkable organ with intrinsic plasticity due to its regenerative and reparative capabilities in response to exercise, traumatic injury, and disease. In Duchenne muscular dystrophy (DMD), the lack of a functional dystrophin protein causes skeletal muscle weakness, dilated cardiomyopathy, respiratory failure, and premature death^1^. Recent progress in the development of *DYSTROPHIN* gene restoration approaches, such as EXONDYS 51 (Eteplirsen), has been shown to benefit a subset of DMD patients amenable to the skipping of *DYSTROPHIN* exon 51 which restores the reading frame^2^. Other *DYSTROPHIN* gene-independent strategies involve the overexpression of a truncated micro-dystrophin (μDYS) construct via adeno-associated viral (AAV) vectors that are under clinical development^3^. The culmination of these strategies centers on improvement of dystrophic pathologies in skeletal and heart muscle, but does not address the systemic effects that occur due to dystrophin protein loss.

A subset of microRNAs (miRNAs) have been identified and shown to be muscle-enriched, play key functional roles in myogenesis and muscle function, and are referred to as myomiRs^4^. In DMD patients, microRNAs isolated from DMD patient serum are utilized as biomarkers and have been referred to as “dystromiRs”^5^. When used in combination with a therapeutic treatment such as exon-skipping phosphorodiamidate morpholino oligomer (PMOs), these dystromiRs can be restored to normal levels making them attractive candidates for marking dystrophic disease progression^6^. miR-486 is a myomiR embedded within the *ANKYRIN1* (*ANK1*) locus and is strictly conserved across mammals^7, 8^. Previously, we demonstrated that miR-486 regulates a DOCK3/PTEN/AKT signaling axis in skeletal muscle, and that muscle-specific overexpression of miR-486 improves dystrophic symptoms in dystrophin-deficient (*mdx^5cv^*) mice^9^. While the reduction of miR-486 expression in dystrophic muscle is known, the suitability of miR-486 as a DMD biomarker, the mechanism for miR-486 dysregulation in DMD muscle, and the functional consequences of the genetic ablation of *miR-486* in mice in normal and dystrophic muscle remains to be elucidated.

We first assessed miR-486 levels in normal and DMD patient skeletal muscles to define its expression with regards to ambulatory status, an important biomarker of disease progression^10, 11^. Expression of miR-486 in ambulatory DMD quadriceps biopsies was significantly lower than healthy quadriceps (**Figure 1A**). miR-486 expression in DMD non-ambulatory muscle also decreased compared to both healthy muscle and ambulatory DMD muscle (**Figure 1A**). Skeletal muscle biopsies from Becker muscular dystrophy (BMD) patients showed decreased miR-486 expression similar to DMD, demonstrating that reduced miR-486 is correlated to pathology related to the loss or decrease of functional dystrophin protein **(Figure 1A).** To evaluate this in a DMD mouse model, we assessed miR-486 levels in three different muscle groups across five DMD disease-relevant time points (1, 3, 6, 9, and 12-months-old) in both *mdx^5cv^* and wild type mice. The *mdx^5cv^* mice demonstrated significantly decreased miR-486 expression in the tibialis anterior (TA), soleus, and diaphragm compared to WT mice and this decrease in expression remained consistent throughout lifespan (**Figure 1B**). MyoD and SRF are important transcriptional activators in skeletal muscle that drive myogenic differentiation of satellite cells into mature myofibers during differentiation and repair^12, 13^. Myogenic differentiation and muscle fiber repair mechanisms are impaired in DMD pathology^14, 15^. We sought to determine if MyoD and SRF signaling played a role in the regulation of miR-486 expression by evaluating MyoD and SRF binding dynamics within the *miR-486* host gene, *ANKRYIN1* (*ANK1*), specifically at the muscle-enriched *ANK1-5* promoter. Using chromatin immunoprecipitation (ChIP), both MyoD and SRF DNA-binding at the human *ANK1-5* promoter were decreased in DMD patient myoblasts and myotubes **(Figure 1C).** This result led us to investigate the expression levels of other MyoD and SRF-related myogenic signaling factors across the lifespan of *mdx^5cv^* mice in affected dystrophic muscles. Quantitative PCR of MyoD and SRF-signaling factors revealed decreased expression in correlation with dystrophic disease progression in the *mdx^5cv^* mouse muscles **(Figure 1D)**.

**Figure 1.**
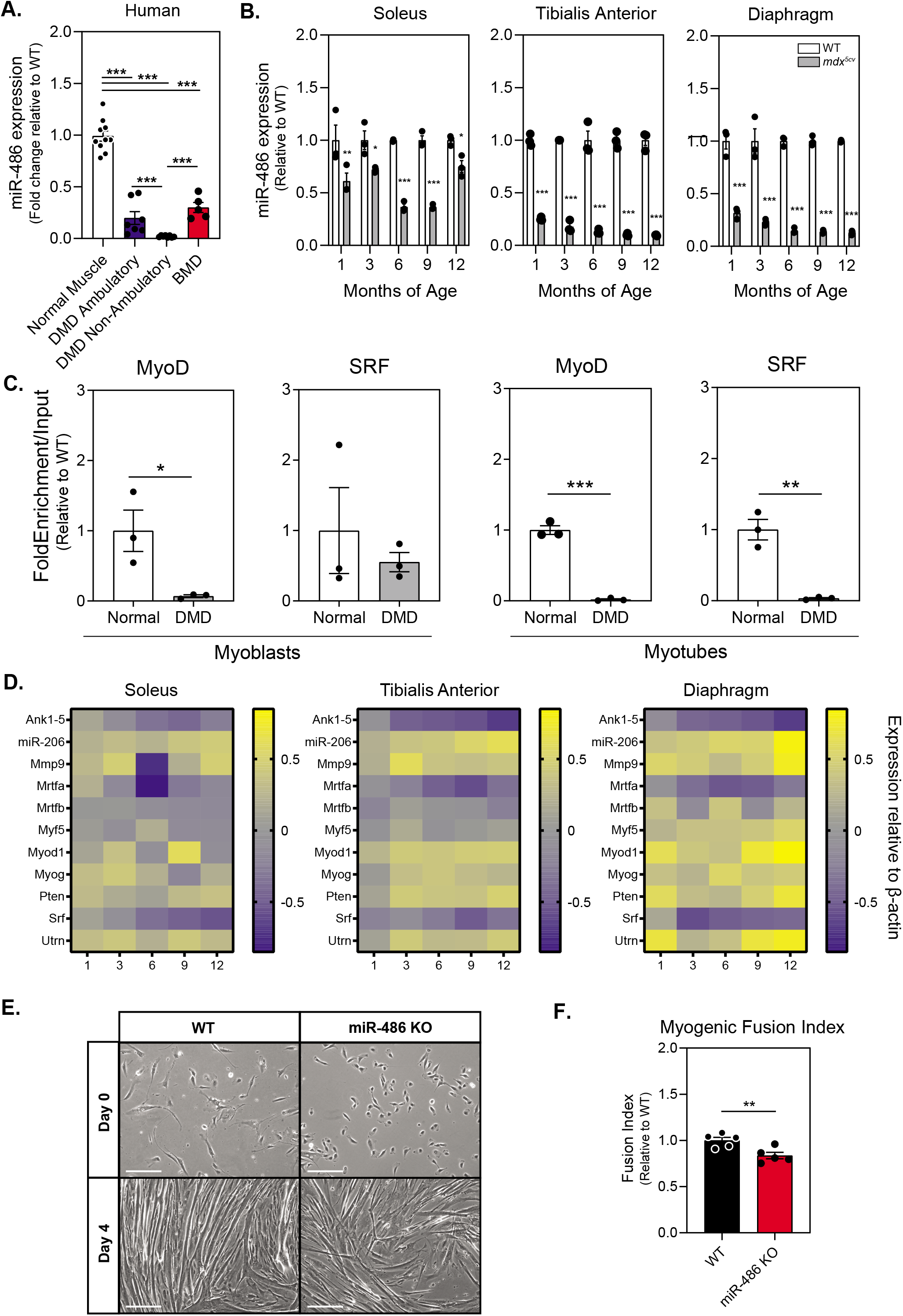
*miR-486* skeletal muscle expression decreases in DMD. **A.** Quantitative PCR reveals decreased expression of miR-486 in dystrophic human skeletal muscle compared to control human skeletal muscle and Becker muscular dystrophy (BMD) muscle. **B.** Quantitative PCR reveals decreasing expression of miR-486 in the *mdx^5cv^* dystrophic mouse model skeletal muscle at 1, 3, 6, 9, and 12 months of age compared to WT control muscle. **C.** ChIP revealed myogenic factors MyoD and SRF demonstrate decreased binding at the promoter of ANK1-5 in isolated *mdx^5cv^* myoblasts and differentiated myotubes compared to WT controls. **D.** Heat maps demonstrate changes in expression of myogenic factors in *mdx^5cv^* tibialis anterior, soleus, and diaphragm muscles over 1, 3, 6, 9, and 12 months of age compared to WT control muscle. Yellow indicates an increase in expression and blue indicates a decrease in expression relative to WT control muscle. **E.** Phase contrast reveals disrupted myoblast fusion and differentiation. Photomicrographs show differentiated myotubes after 4 days of culturing from primary isolated satellite cells. WT and microRNA-486 −/− satellite cells were isolated from <p10 pups and cultured for 4 days. 10X, scale bar = 200μm. **F.** Myogenic fusion index calculated from day 4 myotube culture images depicted in panel **E**. Percentage fusion calculated by dividing the number of nuclei within multinucleated myofibers by the total number of nuclei.

As miR-486 expression is significantly decreased in DMD, we sought to map its functional role in skeletal and cardiac muscle. *miR-486* KO mice generated by CRISPR exhibit an 85 bp deletion within the pre-miRNA hairpin **(Supplemental Figures 1A and 1B)**. Ablation of miR-486 was further validated via Northern blot, as it was undetectable in both *miR-486* KO TA and whole heart **(Supplemental Figure 1C).** As Western blot demonstrated ANK1-5 protein was unperturbed in *miR-486* KO mice, phenotypes in these animals are unlikely to involve disruption of its host gene **(Supplemental Figure 1D).** We evaluated the role of miR-486 in myogenic differentiation by measuring myogenic fusion in the *miR-486 KO* primary myoblasts. The *miR-486* KO myoblasts showed decreased myogenic fusion compared to WT controls (**Figures 1E and 1F**). These results aligned with our previous findings that miR-486 knockdown in myoblasts delayed myogenic differentiation and decreased overall fusion^9^.

To determine how miR-486 loss affects skeletal muscle, we examined histological sections of control and *miR-486 KO* TA muscles (**Figure 2A**). *miR-486 KO* mice exhibit decreased overall myofiber cross-sectional area, increased centralized myonuclei, and increased fibrosis that was exacerbated in the dystrophic *mdx^5cv^* background (**Figures 2B-2D**). We asked if these disruptions played a larger role in whole body composition or locomotion in the *miR-486* KO mice. Interestingly, miR-486 ablation did not significantly alter fat mass, lean mass, or overall total body mass (**Supplemental Figures 2A-C**). As well, *miR-486* KO did not affect extensor digitorum muscle strength, as measured by peak force per unit physiological cross-sectional area (**Supplementa**l **Figure 2D**). DMD-associated dilated cardiomyopathy is a significant cardinal hallmark of advanced DMD pathology^16^. Notably, histopathological analysis of *miR-486* KO mouse hearts showed increased fibrosis compared to WT controls (**Supplemental Figures 3A and 3B**). Echocardiograms of the adult *miR-486* KO mice revealed a decreased fractional shortening, ejection fraction, and several other parameters that were exacerbated in the *miR-486* KO:*mdx^5cv^* (dKO) mice **(Supplemental Figures 3C-3M).** These results demonstrate that miR-486 is involved in pathological remodeling during DMD-associated dilated cardiomyopathy.

**Figure 2.**
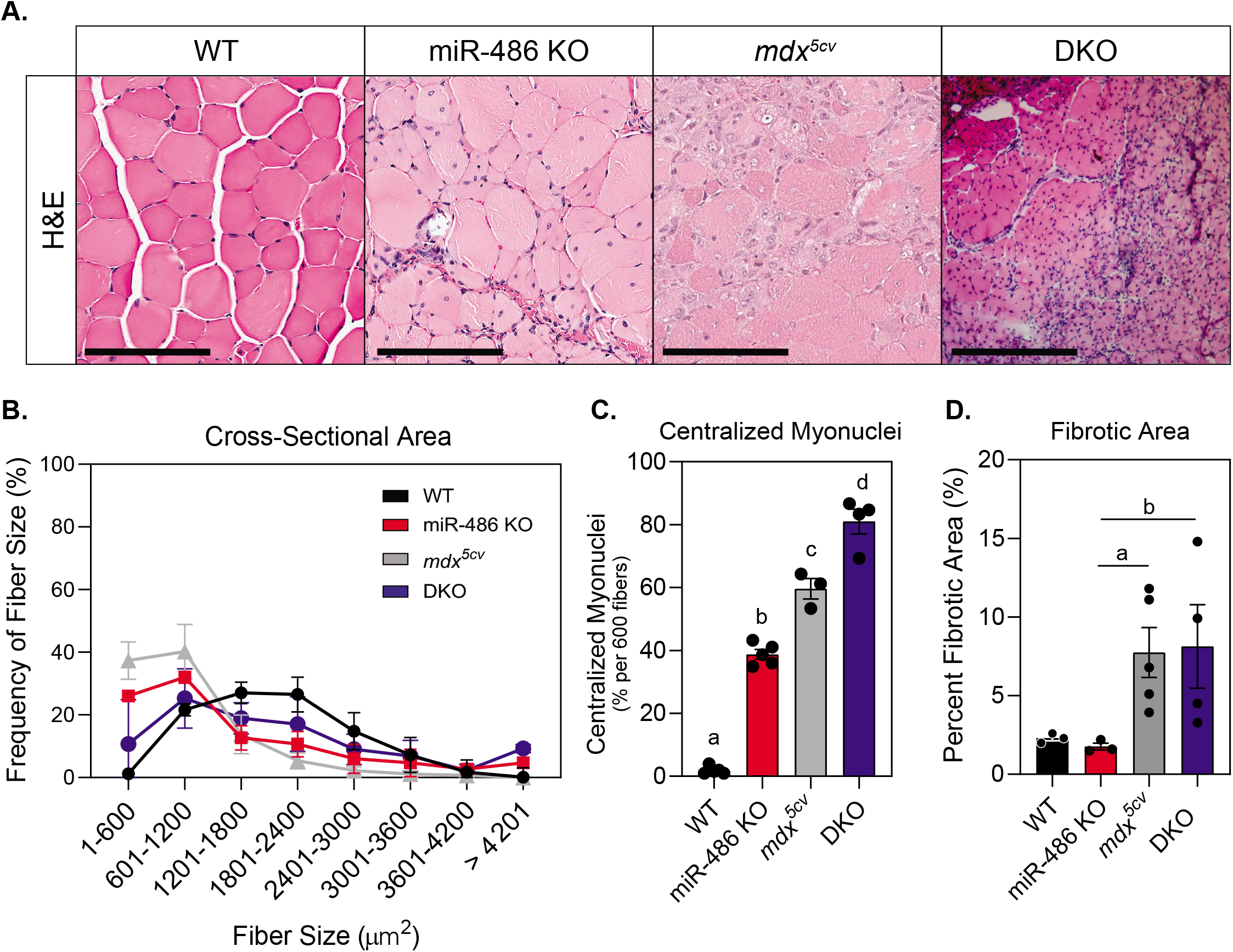
*miR-486* knockout mice demonstrate histological defects in skeletal muscle. **A.** H&E staining of transverse sections of TA muscles at 6 months of age. Scale bars = 200 μM. **B.** Cross-sectional area of myofibers in TA muscles were measured using ImageJ based on H&E staining. 600 fibers from five mice of each genotype were counted. **C.** Centralized myonuclei in WT and *miR-486* KO TA muscle at 6 months of age were counted using ImageJ. 600 fibers from 5 mice of each genotype were counted. Means with different letters are significantly different (Tukey’s HSD, p < 0.05). **D.** Fibrotic area was quantified as a percentage of total area using ImageJ. 5 mice of each genotype were counted. *p=<0.05.

Due to enrichment of miR-486 in skeletal muscle, we sought to define the relationship between the loss of miR-486 and the expression of the four disease-relevant dystromiRs, miR-1, miR-133a, miR-133b and miR-206. Interestingly, only miR-133b was decreased in *miR-486* KO TA muscles **(Figure 3A–D).** To understand the effects of *miR-486* ablation on the overall skeletal muscle transcriptome, we performed RNA sequencing on *miR-486* KO and WT TA muscles. RNA-sequencing revealed 96 transcripts significantly increased in expression and 190 transcripts significantly decreased in expression in *miR-486* KO TA muscles compared to WT **(Figure 3E).** To identify pathways preferentially targeted by miR-486, we utilized g:Profiler software to perform pathway enrichment analysis on the 96 transcripts significantly increased in expression and the 190 transcripts significantly decreased in expression in the *miR-486* KO TA muscles. The top 5 enriched pathways identified by g:Profiler were extracellular matrix, collagen-containing extracellular matrix, regulation of multicellular organismal processes, extracellular region, and vasculature development as shown by the g:GOSt multi-query Manhattan plot **(Figure 3F).** The top 10 transcripts increased in expression and top 10 transcripts decreased in expression in *miR-486* KO compared to WT are shown (**Figure 3G–H**). To evaluate direct targets of miR-486 *in vivo*, we performed chimeric enhanced cross-linking immunoprecipitation-sequencing (eCLIP-seq) to identify miR-486 muscle transcripts that may influence muscle function in complementation with our RNA-seq analysis. Whole TA muscles from WT mice were biopsied and Argonaute-2 (Ago2)-microRNA complexes were immunoprecipitated with miR-486 bound to target mRNA transcripts^17^. MiR-486-specific sequence adaptors were then ligated to the microRNA-mRNA hybrid chimera molecules followed by de-crosslinking and the miR-486-bound transcripts were amplified into a cDNA library (**Figure 4A**). The library was sequenced using *miR-486* KO TA muscles as a control, revealing *miRNA-486* bound targeted transcripts. The chimeric eCLIP-sequencing revealed 18 differentially expressed transcripts that demonstrated reverse complementarity to the miR-486-specific 8-mer (AUGUACUG) binding motif, which overlapped the known conserved miR-486 8-mer seed site (UCCUGUAC) (**Figure 4A**). The eCLIP-seq peak tracks demonstrate the chromosomal location of the 18 differentially expressed transcripts throughout the mouse genome **(Figure 4B).** The sequencing peaks of one of the identified transcripts, *Mt2*, are highlighted using the IGV (Integrated Genome Viewer) sequencing visualization tool **(Figure 4B).** Chimeric eCLIP-seq revealed miR-486 binding to many 3’UTRs but also interestingly to a large numbers of coding sequences (CDS) in muscle genes (**Figures 4C and 4D**). The top 10 peaks that represent mir-486:mRNA bound complexes identified through chimeric eCLIP-seq analysis represent binding sites are listed in **Figure 4E** and the full list of identified peaks is outlined in **Supplemental Data Table 1.** We performed functional enrichment analysis using the g:Profiler tool set to identify key gene ontology (GO) pathways of miR-486 muscle chimeric eCLIP-seq targets^18^. The g:Profiler analysis revealed significant enrichment of miR-486 binding to contractile fiber, myofibril, sarcomere, and other muscle-associated targets as shown by the g:GOSt multi-query Manhattan plot (**Figure 4F**). We next performed quantitative PCR expression analysis on the 18 miR-486 targets (**Supplemental Data Table 1**) identified through eCLIP-seq analysis and compared those to the transcriptome from our *mdx^5cv^* dystrophic mouse TA muscles. The goal was to correlate overlapping DMD biomarkers as many of the miR-486 eCLIP-seq targets had been shown to be dysregulated in DMD patients and involved in key ECM remodeling and metabolic pathways^19^. Unbiased analysis of the 18 miR-486 in vivo targets identified via eCLIP-seq showed that many of them are early stage DMD biomarkers including: *Ttn*, *Camk2a*, *Myom1*, and *Trim63*^20–22^ (**Figures 4E**, **4G**, **Supplemental Data Table 1**). Interestingly, while nearly all of the mRNA transcripts were induced in the *miR-486* KO muscles, several transcripts including those with canonical miR-486 seed sites (*Mt2* and *Auh*) showed variable levels of induction in the absence of dystrophin (**Figure 4G**). The miR-486 targets that are decreased in expression in DMD muscle may be due to the more severe muscle pathologies and overall poor muscle quality observed in the *mdx^5cv^* mice as compared to the *miR-486* KO mice at 6 months of age, as the younger *miR-486* KO mice showed no significant muscle pathological remodeling compared to wild type and *mdx^5cv^* aged-matched cohorts (data not shown). This approach of combining RNA-seq with chimeric eCLIP-seq to identify targets of a key miRNA driving dystrophic disease pathology is useful towards identifying key transcriptomic pathways driving dystrophic muscle pathological remodeling.

**Figure 3.**
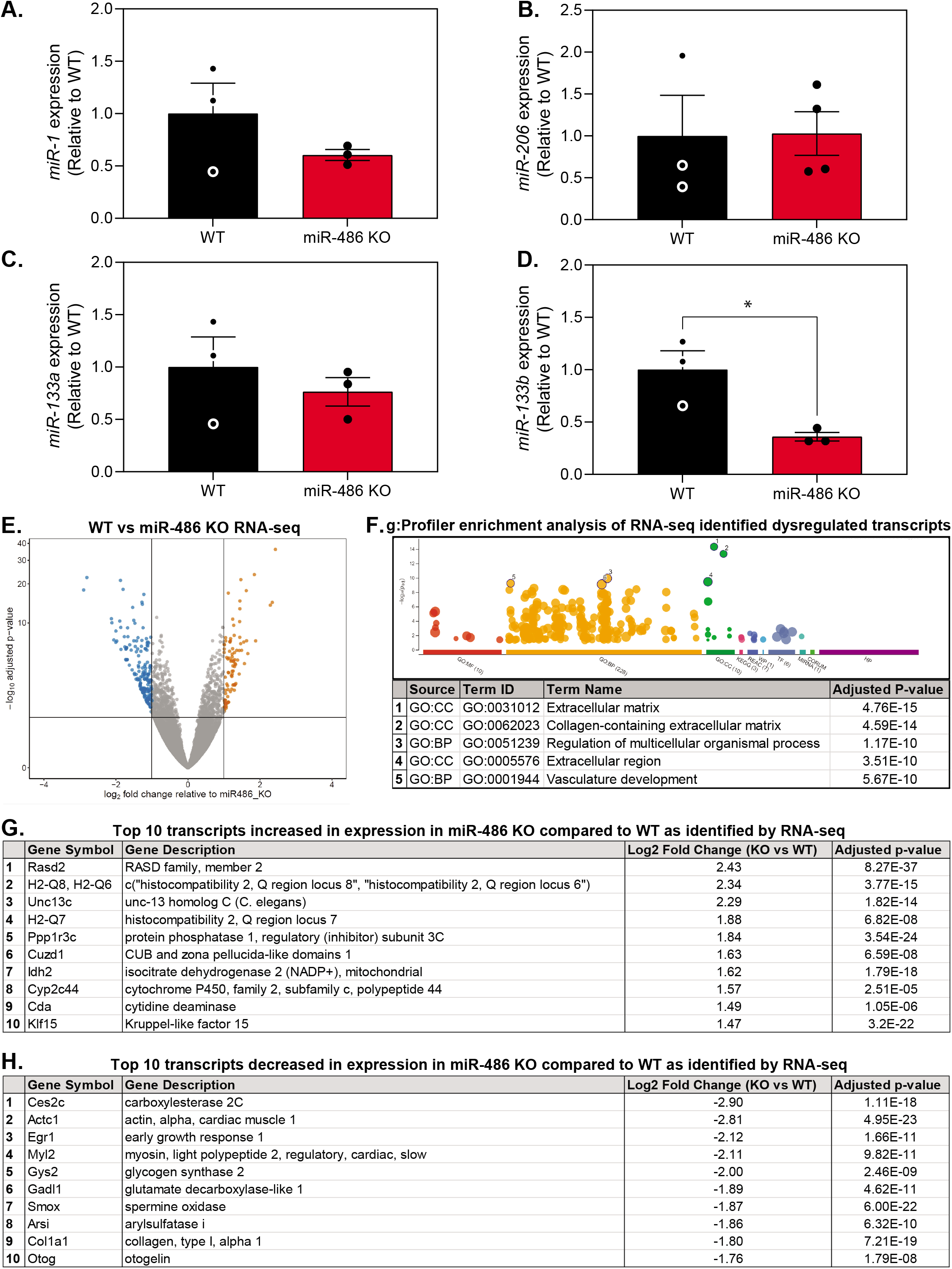
RNA-seq reveals extracellular matrix pathways in *miR-486* KO muscle compared to WT. **A.-D.** Quantitative PCR reveals differential expression of myomiRs in 6-month-old *miR-486* KO TA muscle compared to WT. **E.** Volcano plot demonstrating the fold change and significance of differential gene expression in 6-month-old *miR-486* KO TA muscle compared to WT controls. N=5 mice/cohort were used for comparative analysis. **F.** g:Profiler enrichment analysis of 85 transcripts increased in expression and 159 transcripts decreased in expression based on ≥ 2.0 log_2_ fold change of WT versus *miR-486* KO RNA-seq analysis. The top 5 pathway hits are listed below the graph. **G.** Table of top 10 transcripts increased in expression in 6-month-old miR-486 KO mouse TA muscle compared to WT as identified by RNA-seq. **H.** Table of top 10 transcripts decreased in expression in 6-month-old *miR-486* KO mouse TA muscle compared to WT as identified by RNA-seq.

**Figure 4.**
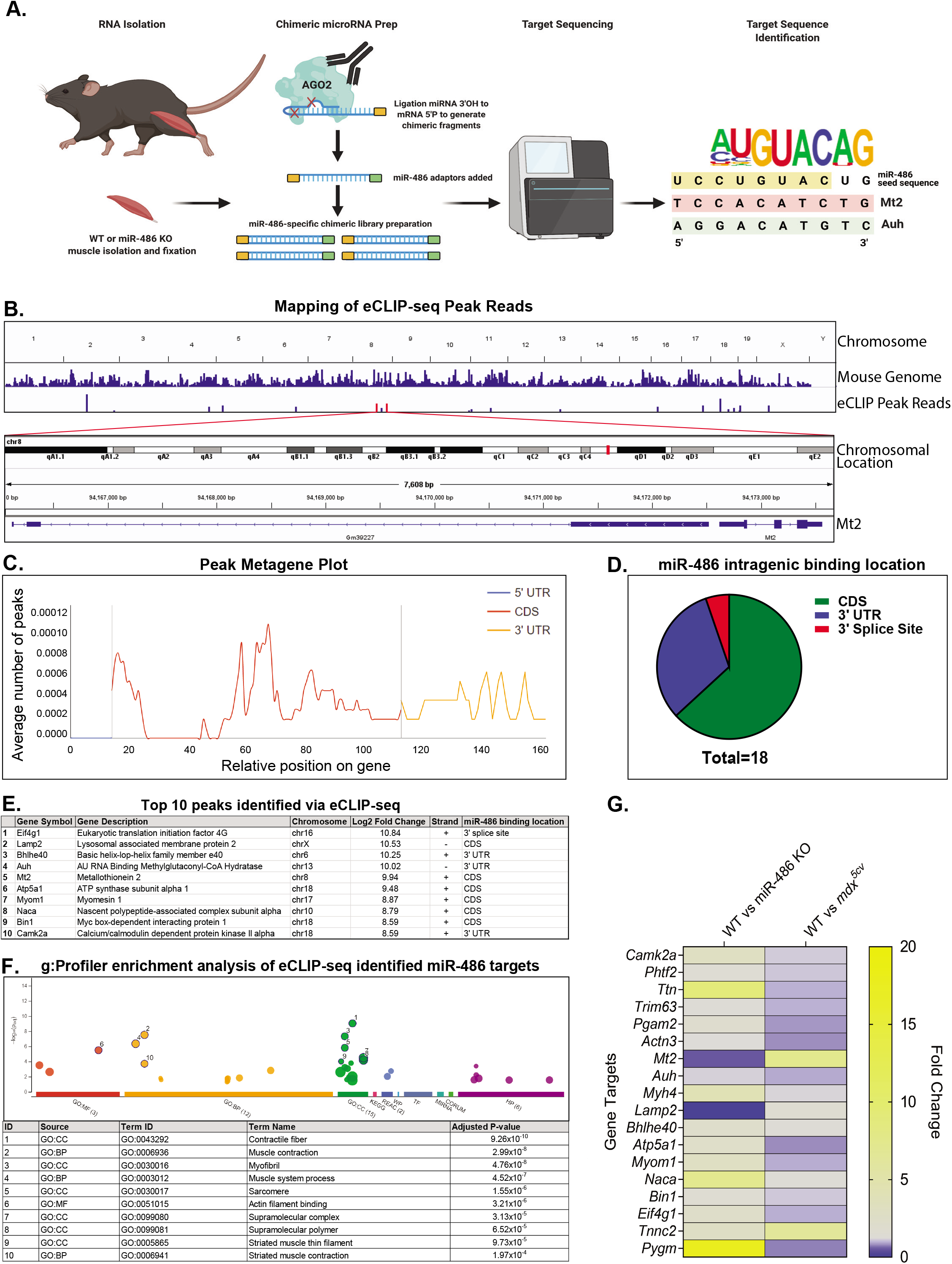
Targeted chimeric eCLIP-seq reveals direct targets of miR-486. **A.** Schematic demonstrating the workflow for the chimeric eCLIP-sequencing platform to identify miR-486 *in vivo* skeletal muscle target transcripts. TA muscle was harvested from six-month-old WT and *miR-486 KO* male mice and total RNA isolation was completed. The Ago2-miR-486 complex bound to target RNA transcripts was isolated and then sequencing was performed to map the reads of transcripts bound to the Ago2-miR-486 complex. **B.** Chromosomal location of a single top “hit”, Mt2, as identified by eCLIP-seq as a direct target of miR-486. Peaks generated using Integrative Genome Viewer. **C.** Metagene plot demonstrating overall miR-486 binding location by relative position on target gene. **D.** Pie chart demonstrating the proportion of miR-486 gene targets and the respective intragenic binding location of miR-486. **E.** Table outlining the top 10 transcripts that were identified as direct targets of miR-486 via CLIP-seq. **F.** g:Profiler enrichment analysis graph demonstrates the most significant cellular pathways associated with the 18 direct miR-486 targets identified via chimeric eCLIP-seq. The pathway ID number in the table correlates with the numbered dots in the accompanying graph above. **G.** Quantitative PCR of 18 miR-486 eCLIP-seq targets in miR-486 KO and *mdx^5cv^* TA muscles expression levels compared to WT controls in separate cohort analyses. Data points are individual biological replicates, n=4/cohort and logarithmic fold change normalized to both wild type and β-actin is shown. **p=≤0.01. mRNA levels are normalized to β-actin and miR-486 KO levels are shown as relative to WT.

The dynamic nature of miR-486 expression and its role in both pathological muscle remodeling and metabolic regulation suggests its requirement in muscle during times of stress and disease. Our findings demonstrate that miR-486 is a key DMD biomarker whose downregulation drives pathogenic remodeling of dystrophic muscle by targeting key muscle structural, metabolic, and extracellular matrix remodeling factors. Indeed, miR-486 is required for normal skeletal and cardiac muscle growth and function through modulation of these factors as well. Loss of miR-486 expression has profound consequences on mouse skeletal and cardiac muscle architecture, fibrotic depositions, centralized myonuclei, and overall muscle function. In cardiac muscle, miR-486 ablation resulted in significant pathological remodeling of the heart and overall decreased cardiac output. There is growing evidence that restoration of dystrophin via exon-skipping compounds can restore the transcriptome signature of dystrophic muscle, including miRNAs, to that of normal or Becker-like muscle^23^. Our combinatorial RNA-sequencing and eCLIP-seq of skeletal muscles from miR-486 KO mice identified a network of miR-486 muscle targets including coding and non-coding regions of muscle targets that play key roles in dystrophin-deficient muscle pathology. MicroRNA profiling studies demonstrated that miR-486 levels increased in the right ventricles of hypoplastic left heart syndrome (HLHS) patients, and miR-486 is elevated in response to cyclic stretch^24^. Other studies demonstrate that miR-486 levels are modulated during development, stress, and disease states via modulation of key cellular growth pathways through direct regulation of FOXO, SMAD, PI3K, and AKT^8, 9, 24, 25^. These growth pathways are also dysregulated in dystrophic skeletal muscles and can be targeted therapeutically via miR-486 transgenic overexpression in skeletal muscle to restore proper signaling pathways^8, 9^.

Additional questions remain as to which specific myogenic cell populations and during which stages of dystrophic development miR-486 dysregulation occurs. Several studies have identified miR-486 enrichment in quiescent muscle satellite cells, along with other muscle-enriched myomiRs prior to activation^26^. As activation of muscle satellite cells occurs along with myoblast fusion, miR-486 expression increases which may represent the switch from MyoD-driven miR-486 expression to SRF-driven expression that we show at the *miR-486/ANK1-5* locus. While miR-486 expression was shown to regulate Pax7 in quiescent mouse muscle satellite cells, there are likely additional miR-486 targets in MSCs as well as other non-muscle cell types. Furthermore, many of the miR-486 mRNA targets we identified are also expressed in non-muscle cell types. This suggests that additional levels of post-transcriptional regulation may yield novel tissue-specific miR-486 target transcripts.

MicroRNA-486 overexpression may be a valid means of restoration of key dysregulated signaling pathways in dystrophic muscle that affect growth and muscle function. There is mounting evidence that the collagens and the ECM remodeling that occurs in DMD serves as a biomarker for disease progression and a therapeutic target^27–29^. Additional evidence demonstrating the restoration of microRNAs in exon-skipped *mdx* mice, suggests that miR-486 expression profiling is highly useful in evaluating dystrophic replacement and restoration strategies in muscle^6, 30–32^. Given our previous work demonstrating that miR-486 transgenic overexpression in a dystrophic mouse model can ameliorate dystrophic pathologies, strategies to either induce miR-486 expression or direct miR-486 overexpression should be explored for therapeutic disease modulation.

## Acknowledgements

The authors wish to thank Alexander Shishkin, Kylie Shen, Sergei Manakov, Heather Foster, and Gene Yeo from Eclipse BioInnovations for their performance and assistance with the chimeric eCLIP-sequencing experiments. We wish to thank Margaret Bell, Kaleen Lavin, Layne Peacock, and Gina Seay for assistance with the collection of normal human muscle control biopsies. We wish to thank Emanuela Gussoni and Glenn Rowe for a critical reading of the manuscript.

## Author Contributions

R.M.H., A.S., A.L.R., K.G.E., M.A.L., D.A.S., M.M.B, G.C.R, J.J.W., D.K.C., E.C.L., M.X, and M.S.A. all performed experiments or analysis of data. M.A.L., M.M.B., M.J.C., J.S.D., D.J., E.C.L., M.X., and M.S.A. provided unique reagents. R.M.H., A.S., A.L.R., and M.S.A. analyzed all the data and wrote the manuscript. All authors approved of the manuscript prior to submission.

## Funding Agencies

R.M.H. is a funded by an NIH National Institute of Neurological Disorders and Stroke (NINDS) F99/K00 grant (F99NS118718). M.A.L. is funded by an NIH NINDS K08 grant NS120812 and a grant from the Kaul Pediatric Research Institute. The UAB Small Animal Phenotyping Core supported by the NIH Nutrition & Obesity Research Center (P30DK056336), and the Mouse Cardiovascular Core Vevo 3100 Mouse Ultrasound Facility for this project. Research reported in this publication was supported by Eunice Kennedy Shriver National Institute of Child Health and Human Development, NIH, HHS of the National Institutes of Health under award number R01HD095897 awarded to M.S.A. M.S.A. is also a co-investigator on an NIH NIAMS award R21AR074006 and is also funded by a Muscular Dystrophy Association (MDA) grant (MDA41854). M.M.B. was supported by NIH NIAMS U01AR071133 and NICHD P2HD086851 grants. M.X. was supported by NIH NHLBI grants R01HL153501 and R03HL141620. Work in the E.C.L. lab was supported by NIH grants R01GM083300 and R01HL135564, and by the MSK Core Grant P30CA008748.

## List of Abbreviations

Ago2: Argonaute-2
ANK1: Ankyrin1
BMD: Becker muscular dystrophy
CDS: coding sequence
ChIP: chromatin immunoprecipitation
eCLIP: enhanced cross-linking and immunoprecipitation
CSA: cross-sectional area
DAPC: Dystrophin-associated complex
DEXA: dual energy x-ray absorptiometry
DMD: Duchenne muscular dystrophy
ECM: extracellular matrix
GTT: glucose tolerance test
IGV: Integrated Genome Viewer
MyHC: myosin heavy chain
MSC: muscle satellite cell
PMO: phosphorodiamidate morpholino oligomer
QMR: Quantitative Magnetic Resonance
qPCR: quantitative polymerase chain reaction
sANK1: small Ankryin1/Ank1-5
Seq: sequencing
TA: Tibialis anterior

## Methods

### Mice

The *miR-486* KO mice were generated as previously described^33^. Mice were initially maintained on the *FVB/NJ* (Jackson Labs; #001800) strain and were later backcrossed onto the *C57BL/6J* (Jackson Labs; #000664) strain for more than six generations to ensure isogenicity. All analyses were performed on the same strain background. The *mdx^5cv^* (Jackson Labs; #002379) were also kept on the *C57BL/6J* strain, and homozygote mutant females were mated to the *miR-486* KO males to generate *miR-486* +/−:*mdx^5cv^* mice and later *miR-486* KO:*mdx^5cv^* (DKO) mice. Unless otherwise stated, only 6-month-old male mice were used in all experimental conditions. All mouse strains were maintained under standard housing and feeding conditions with the University of Alabama at Birmingham Animal Resources Facility under pathogen-free conditions and all protocols were approved under the animal protocol number 21393.

### Chromatin Immunoprecipitation (ChIP)

Chromatin immunoprecipitation was performed on normal human and DMD myoblasts and myotubes in a protocol previously described^34^. Genomic DNA was isolated from myoblast (70% confluency in growth medium) and myotube (4 days differentiation in low serum differentiation medium) and ChIP was performed using the Simple ChIP Enzymatic Chromatin IP Kit (Cat# 9003; Cell Signaling Technology; Danvers, MA) following the manufacturer’s protocols. For the MyoD ChIP assays, a rabbit polyclonal (MyoD clone M-318; Cat# sc-760; Santa Cruz Biotechnology; Dallas, TX) was used to immunoprecipitate MyoD bound to the mouse *miR-486* locus. For SRF ChIP assays, a rabbit polyclonal (SRF clone G-20; Cat# sc-335; Santa Cruz Biotechnology; Dallas, TX) was used to immunoprecipitate SRF bound to the mouse *miR-486* locus. Myoblasts were seeded at 4 ×10^5^ cells/15 cm dish in 10 dishes per cohort. The myoblast fraction consisted of normal human myoblasts grown to approximately 70% confluency. For the myotube fraction (five 15 cm dishes), after reaching 90% confluency, myoblasts were differentiated into myotubes by serum withdrawal (2% FBS) for 4 days. Cells were harvested following three washes in ice-cold 1 X DPBS, were crosslinked with 1% formaldehyde (Cat# F8775; Sigma-Aldrich; St. Louis, MO) for one hour at room temperature, and then chromatin was sheared using by incubating the DNA pellets with micrococcal nuclease for 20 min at 37°C. Ten percent of chromatin was removed for input controls, while the approximately 200 mg of total chromatin was used to purify DNA fragments following immunoprecipitation with 5 mg of either MyoD or SRF antibodies. The following morning, the samples were incubated with 30 ml of ChIP Grade Protein G magnetic beads (Cat# 9006S; Cell Signaling Technology; Danvers, MA) for 2 hours at 4°C under gentle rotation. Following several stringent washes, the chromatin was eluted off the columns in 1 mM Tris-HCL pH 8.0 (Cat# 15568025; ThermoFisher; Waltham, MA) and purified on DNA-binding columns (Cat# D4029; Zymo Research; Tustin, CA) before a final elution in sterile water. MyoD and SRF ChIP-seq annotation of ANK1-5 was completed using the publicly available UCSC Genome Browser track data hubs annotation resource available at genome.ucsc.edu. For the MyoD TF binding site at the human *ANK1-5* locus the following primers were used Fwd: 5’-GAGCTCCAAGACTGAGGACTGGAC-3’ and Rev: 5’-CAGGGAGGATGGAGATCAGAGCC-3’. The non-specific MyoD site negative control primers next to the human *ANK1-5* locus used were Fwd: 5’- CTGCACGTCAGCCTCCCAAAG-3’ and Rev: 5’- ACTGGGATCCTCCAGGGGCC-3’. The MyoD TF positive control primers used were Fwd: 5’-CAGTGAACAATGGTGCTTGG-3’ and Rev: 5’- TTCCACATTCACGCAGAGAG-3’. For the SRF TF binding site at the human *ANK1-5* locus the following primers were used Fwd: 5’-ACAGTAGGTGAGTTGCAGGGTTAG-3’ and Rev: 5’-GGGCTCAGGGACAGTCAAGTGAGC-3’. The non-specific SRF site negative control primers next to the human *ANK1-5* locus used were Fwd: 5’-GAAACACGGAGCAGCCTGGC-3’ and Rev: 5’-ATGAGGATCGACTGTACATGC-3’. The SRF TF positive control primers used were Fwd: 5’- TGGTTGGATAACAGAGGCAGA-3’ and Rev: 5’-GCTTCTGTTGTGGCGTCTTT-3’.

### Western blots

Protein lysates from cell and tissues were lysed and homogenized (tissues) in Mammalian Protein Extraction Reagent (M-Per) lysis buffer (Cat# 78501; ThermoFisher; Waltham, MA) supplemented with cOmplete protease inhibitor tablets (Cat# 1183617000; MilliporeSigma; Burlington, MA). Approximately 50 μg of lysate were electrophetically resolved on 4-20% Novex Tris-glycine gradient gels (Cat# XV04200PK20; ThermoFisher; Waltham, MA). Proteins were then transferred to 0.2 μm PVDF membranes (Cat# 88520; ThermoFisher; Waltham, MA), and incubated overnight at 4°C with gentle rocking in primary antisera diluted 1:1000 in 5% bovine serum albumin (Cat# A30075; RPI Corp; Mount Prospect, IL)/1xTBS-Tween (Cat#IBB-581X; BostonBioProducts; Ashland, MA). Membranes were washed three times in 1xTBS-Tween for 5 minutes each, and then incubated in secondary antisera (1:2000 dilution) for 1 hour at room temperature. Membranes were then washed four times in 1x TBS-Tween for 15 minutes each before the addition of Novex Chemoluminescent Substrate Reagent Kit (Cat# WP20005; ThermoFisher; Waltham, MA). Membranes were exposed onto PerfectFilm audioradiography film (Cat#B581; GenHunter; Nashville, TN) and developed on a Typhoon Variable Mode Imager (Amersham Pharmacia; Little Chalfont, United Kingdom). Western blot densitometry was performed using open-source ImageJ software.

#### Immunofluorescence of Muscle Sections

For immunofluorescent staining, frozen sections were fixed in 100% acetone for 5 minutes at room temperature, then air dried for 20 minutes. The slides were then incubated was in 1x PBS-Tween (Cat#IBB-171R; BostonBioProducts; Ashland, MA). To reduce non-specific binding, the slides were then incubated with blocking reagent from the Mouse-on-Mouse (M.O.M) kit (Cat# BMK-2202; Vector Laboratories; Burlingame, CA). Slides were incubated for one hour at room temperature in primary antibody. Slides were imaged with a Zeiss LSM 700 Laser Scanning Confocal at ×10 objective, and Zen software (Carl Zeiss Microscopy, LLC) was used for image capture. ImageJ software was used to generate montages of muscle sections for fiber counting. Adobe Photoshop version 2020 (Adobe Systems Inc.; San Jose, CA) was used to adjust resolution and contrast for representative images.

#### Muscle Histochemical Staining

Mouse skeletal muscle tissues were cryo-frozen by covering the tissue Optimum Cutting Temperature (OCT) solution (FisherScientific; Hampton, NH; Cat#23-730-571) and completely submerging the tissues in a liquid nitrogen-chilled Isopentane (Cat#AC397221000; FisherScientific; Hampton, NH) bath as unfixed tissues. Blocks were later cut on a cryostat and 7-10 μm thick sections were placed on SuperFrost Plus slides (Cat#FT4981GLPLUS; ThermoFisher; Waltham, MA). Hematoxylin and eosin (H&E) staining was performed as previously described^8^. Mouse skeletal muscle tissues were cryo-frozen and sectioned as described above. Masson’s Trichrome staining was performed on frozen sections using the Masson’s Trichrome stain kit (Cat# 25088-100; Polysciences; Warrington, PA) and the following the manufacturer’s protocol. Mouse hearts were perfusion fixed in 10% neutral buffered formalin (Cat#HT501128; MilliporeSigma; Burlington, MA) overnight at 4°C and held in 70% ethanol for 24-48 hours before being processed as previously described^35^.

#### RNA-sequencing and data analyses

Adult 6-month-old male TA muscles from WT and *miR-486* KO mice (n = 4 mice per cohort) were snap frozen in liquid nitrogen. Muscle samples were mechanically homogenized and total RNA was extracted using a miRVana Isolation Kit (Cat#AM1560; ThermoFisher; Waltham, MA) following the manufacturer’s protocol. The total RNA was amplified using the Sure Select Stranded RNA-Seq kit (Agilent Technologies; Santa Clara, CA) using standard protocols. A ribominus kit (Cat# K155002; ThermoFisher; Waltham, MA) was used to deplete large ribosomal RNAs. All biological replicates contained a minimum of 35.7 million reads with an average number of 39.6 million reads across the replicates. The FASTQ files were uploaded to the UAB High Performance Computer cluster for bioinformatics analysis with the following custom pipeline built in the Snakemake workflow system (v5.2.2)^36^: first, quality and control of the reads were assessed using FastQC, and trimming of the Illumina adapters and bases with quality scores of less than 20 were performed with Trim_Galore! (v0.4.5). Following trimming, the transcripts were quasi-mapped and quantified with Salmon^37^ (v0.12.0, with ‘--gencode’ flag for index generation and ‘-l ISR’, ‘--gcBias’ and ‘--validateMappings’ flags for quasi-mapping) to the mm10 mouse transcriptome from Gencode release 21. The average quasi-mapping rate was 70.4% and the logs of reports were summarized and visualized using MultiQC^38^ (v1.6). The quantification results were imported into a local RStudio session (R version 3.5.3) and the package “tximport”^39^ (v1.10.0) was utilized for gene-level summarization. Differential expression analysis was conducted with DESeq2^40^ package (v1.22.1). Following count normalization, principal component analysis (PCA) was performed, and genes were defined as differentially expressed genes (DEGs) if they passed a statistical cutoff containing an adjusted p-value <0.05 (Benjamini-Hochberg False Discovery Rate (FDR) method) and if they contained an absolute log2 fold change >=1. Functional annotation enrichment analysis was performed in the NIH Database for Annotation, Visualization and Integrated Discovery (DAVID, v6.8) by separately submitting upregulated and downregulated DEGs. A p-value <0.05 cutoff was applied to identify gene ontology (GO) terms. The FASTQ files of the current study have been uploaded to NCBI’s Gene Expression Omnibus under accession number GSE155787.

#### Chimeric eCLIP-Sequencing

For the chimeric CLIP-sequencing we used adult 6-month-old male TA muscles from WT and *miR-486* KO mice (n = 3 muscles for each cohort) that were snap frozen in liquid nitrogen and chimeric eCLIP-sequencing was performed with assistance by Eclipse BioInnovations (Eclipse BioInnovations; San Diego, CA). The FASTQ files of the current study have been uploaded to NCBI’s Gene Expression Omnibus under accession number GSE173821. A detailed description of the chimeric CLIP-seq procedures including Ago2 immunoprecipitation, library amplification, and other quality control analyses can be found in the **Supplemental Figures and Methods.**

#### Real time quantitative PCR (rt-qPCR)

Total RNA was extracted from muscle tissue using the miRVana (Cat# AM1560; ThermoFisher; Waltham, MA) kit following the manufacturer’s protocol. One microgram of total RNA was reverse transcribed using a Taqman Reverse Transcription kit following the manufacturer’s protocol (Cat# N8080234; Applied Biosystems; Foster City, CA). For microRNA fractions, 50 nanograms of total small RNA was used for all reverse transcription reactions. Taqman assay probes were purchased from Applied Biosystems corresponding to the individual genes. Quantitative PCR (qPCR) Taqman reactions were performed using Taqman Universal PCR Master Mix (Applied Biosystems; Cat# 4304437). Samples were run on the Fluidigm Biomark HD system (Fluidigm Corp.; San Francisco, CA) in 96.96 Dynamic Array plates. Relative expression values were calculated using the 2^−ΔΔCt^ method.

