## Supplemental figures for "miR-486 is an epigenetic modulator of Duchenne muscular dystrophy pathologies"

Supplemental Figure 1.

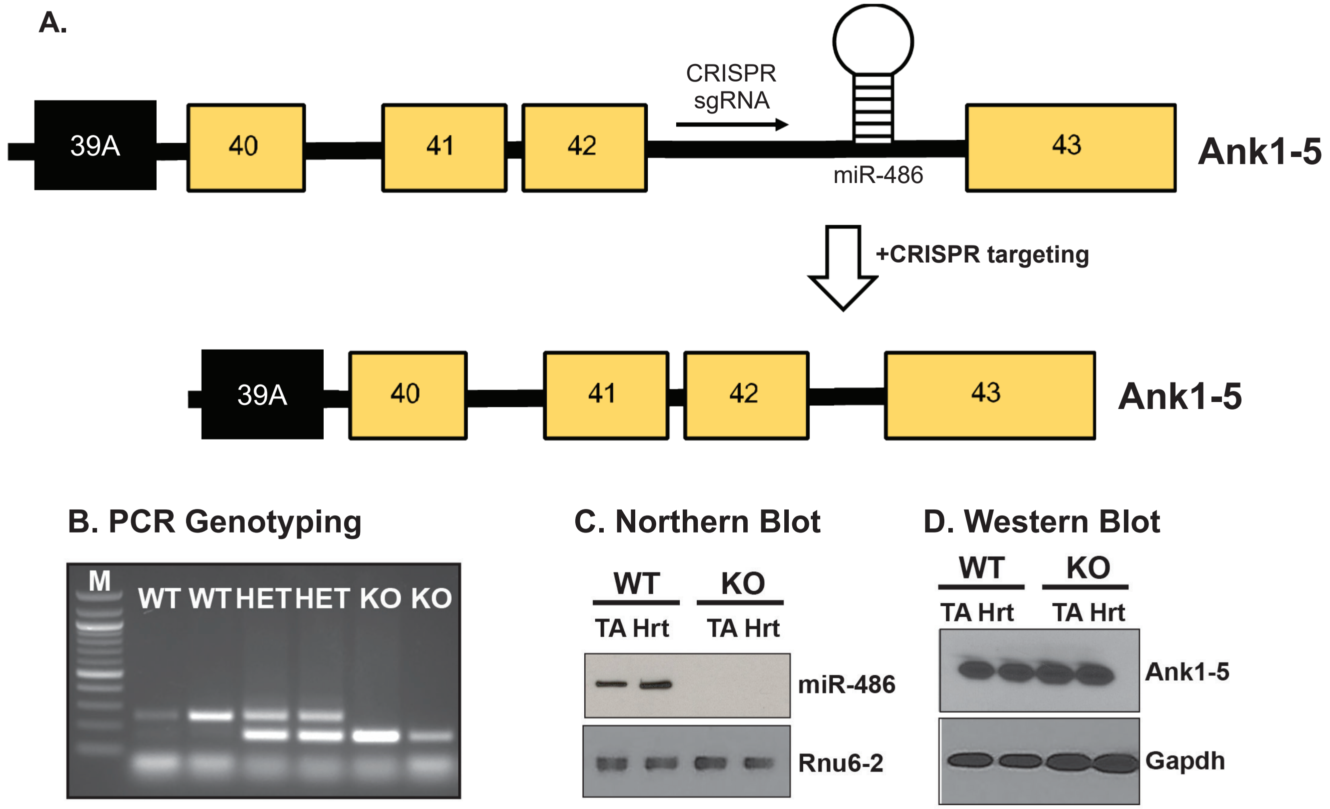

**Supplemental Figure 2.**

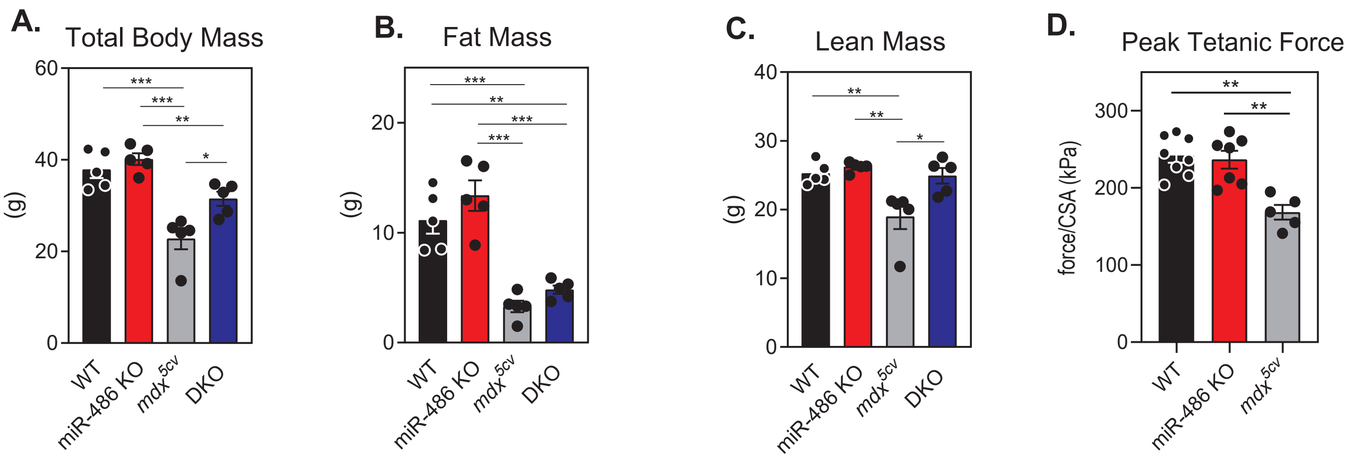

**Supplemental Figure 3.**

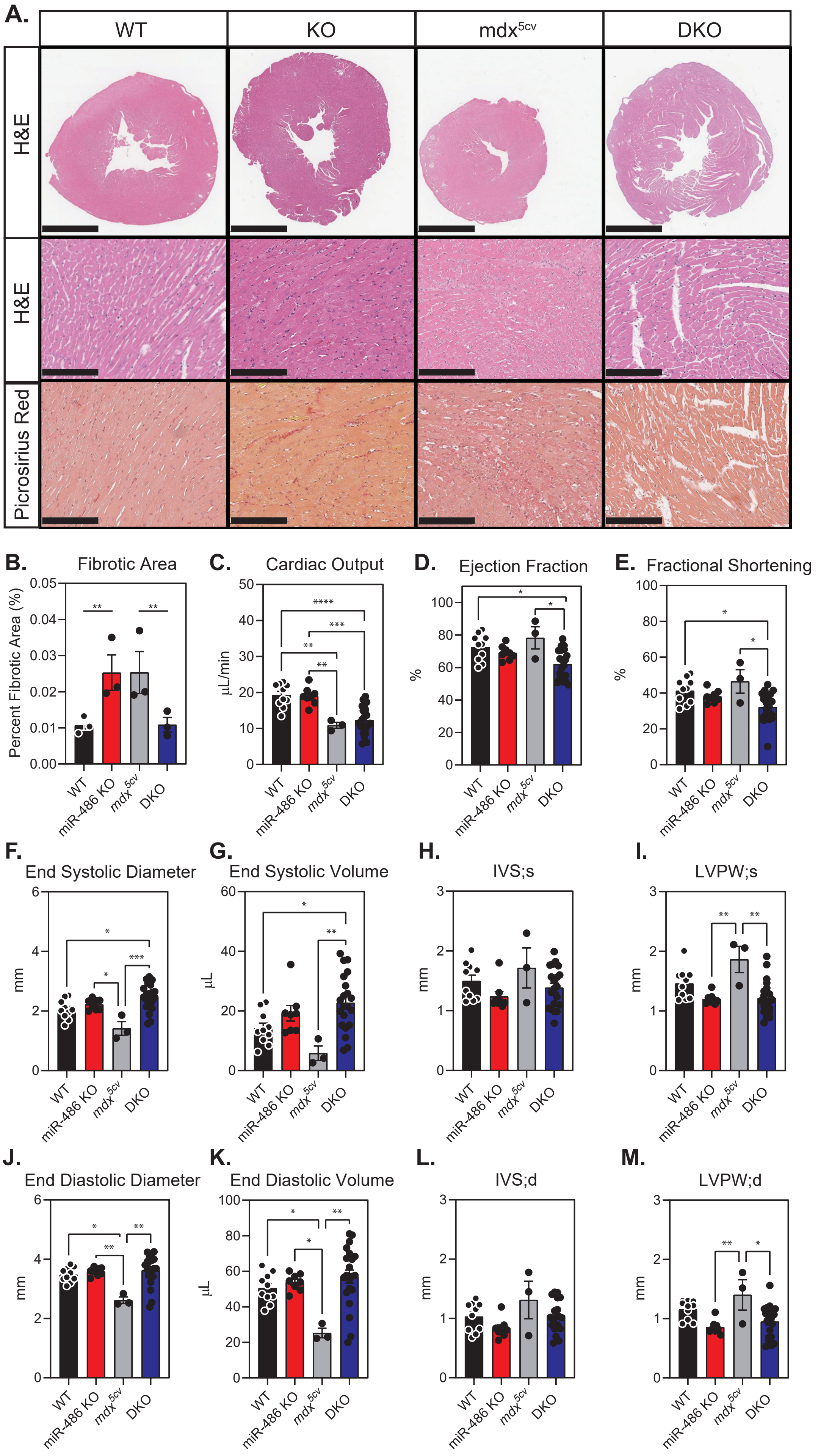

### Supplemental Data Table 1

#### Full list of peaks identified via CLIP-seq

| Chromosome | Log2 Fold Change | Strand | Gene Symbol | Gene Name | miR-486 binding location |
|---|---|---|---|---|---|
| chr16 | 10.83631252 | + | Eif4g1 | Eukaryotic translation initiation factor 4G | 3' splice site |
| chrX | 10.5286901 | - | Lamp2 | Lysosomal associated membrane protein 2 | CDS |
| chr6 | 10.24505513 | + | Bhlhe40 | Basic helix-loop-helix family member e40 | 3' UTR |
| chr13 | 10.02281843 | - | Auh | AU RNA binding methylglutaconyl-CoA Hydratase | 3' UTR |
| chr8 | 9.935358976 | + | Mt2 | Metallothionein 2 | CDS |
| chr18 | 9.481085019 | + | Atp5a1 | AtP synthase subunit alpha 1 | CDS |
| chr17 | 8.86887411 | + | Myom1 | Myomesin 1 | CDS |
| chr10 | 8.794192464 | + | Naca | Nascent polypeptide-associated complex subunit alpha | CDS |
| chr18 | 8.589497725 | + | Bin1 | Myc box-dependent interacting protein 1 | CDS |
| chr18 | 8.589283851 | + | Camk2a | Calcium/calmodulin dependent protein kinase II alpha | 3' UTR |
| chr5 | 8.521814738 | - | Phtf2 | Putative homeodomain transcription factor 2 | 3' UTR |
| chr2 | 8.521577645 | - | Ttn | Titin | CDS |
| chr4 | 8.393954899 | + | Trim63 | Tripartite motif containing 63 | 3' UTR |
| chr19 | 8.299717786 | + | Pygm | Glycogen phosphorylase muscle associated | CDS |
| chr11 | 6.623766942 | - | Pgam2 | Phosphoglycerate mutase 2 | CDS |
| chr11 | 5.663698166 | + | Myh4 | Myosin heavy chain 4 | CDS |
| chr19 | 5.567377114 | - | Actn3 | Alpha-actinin-3 | CDS |
| chr2 | 4.791205601 | - | Tnnc2 | Troponin C2 | CDS |
